# PARP inhibition and pharmacological ascorbate demonstrate synergy in castration-resistant prostate cancer

**DOI:** 10.1101/2023.03.23.533944

**Authors:** Nicolas Gordon, Peter T. Gallagher, Neermala Poudel Neupane, Amy C. Mandigo, Jennifer K. McCann, Emanuela Dylgjeri, Irina Vasilevskaya, Christopher McNair, Channing J. Paller, Wm. Kevin Kelly, Karen E. Knudsen, Ayesha A. Shafi, Matthew J. Schiewer

## Abstract

Prostate cancer (PCa) is the second leading cause of cancer death for men in the United States. While organ-confined disease has reasonable expectation of cure, metastatic PCa is universally fatal upon recurrence during hormone therapy, a stage termed castration-resistant prostate cancer (CRPC). Until such time as molecularly defined subtypes can be identified and targeted using precision medicine, it is necessary to investigate new therapies that may apply to the CRPC population as a whole.

The administration of ascorbate, more commonly known as ascorbic acid or Vitamin C, has proved lethal to and highly selective for a variety of cancer cell types. There are several mechanisms currently under investigation to explain how ascorbate exerts anti-cancer effects. A simplified model depicts ascorbate as a pro-drug for reactive oxygen species (ROS), which accumulate intracellularly and generate DNA damage. It was therefore hypothesized that poly(ADP-ribose) polymerase (PARP) inhibitors, by inhibiting DNA damage repair, would augment the toxicity of ascorbate.

**Results:** Two distinct CRPC models were found to be sensitive to physiologically relevant doses of ascorbate. Moreover, additional studies indicate that ascorbate inhibits CRPC growth *in vitro* via multiple mechanisms including disruption of cellular energy dynamics and accumulation of DNA damage. Combination studies were performed in CRPC models with ascorbate in conjunction with escalating doses of three different PARP inhibitors (niraparib, olaparib, and talazoparib). The addition of ascorbate augmented the toxicity of all three PARP inhibitors and proved synergistic with olaparib in both CRPC models. Finally, the combination of olaparib and ascorbate was tested *in vivo* in both castrated and non-castrated models. In both cohorts, the combination treatment significantly delayed tumor growth compared to monotherapy or untreated control.

**Conclusions:** These data indicate that pharmacological ascorbate is an effective monotherapy at physiological concentrations and kills CRPC cells. Ascorbate-induced tumor cell death was associated with disruption of cellular energy dynamics and accumulation of DNA damage. The addition of PARP inhibition increased the extent of DNA damage and proved effective at slowing CRPC growth both *in vitro* and *in vivo*. These findings nominate ascorbate and PARPi as a novel therapeutic regimen that has the potential to improve CRPC patient outcomes.

## Introduction

The American Cancer Society projects 288,300 new cases of prostate cancer (PCa) will be diagnosed and estimates that PCa will result in 34,700 deaths in 2023^1^. Most prostate cancer-related deaths occur due to metastatic dissemination. Metastatic PCa responds to androgen deprivation therapy (ADT), sometimes referred to as chemical or medical castration; however, the cancer often recurs within 2-3 years^2^. At this point, the disease is termed metastatic castrationresistant prostate cancer (mCRPC). Treatment options for patients with mCRPC are limited to more recently developed anti-androgen agents such as enzalutamide and abiraterone, as well as radium-223, docetaxel, and cabazitaxel; however, these agents are typically not curative^3,4^. In short, the mCRPC population is in dire need of new therapeutic options.

The use of ascorbate, more commonly known as Vitamin C, in cancer treatment has been controversial for decades. In the 1970’s, Linus Pauling and Ewan Cameron demonstrated survival benefit for patients treated with ascorbate in a variety of end-stage cancers^5^. Subsequent studies failed to show any benefit however, and the concept of ascorbate as an anti-cancer agent was abandoned^6^. More recent research has revealed significantly limited bioavailability for orally administered ascorbate which explains why Cameron and Pauling’s results with intravenous ascorbate were not replicated in the consequent studies designed to assess the benefits of oral supplementation. Intravenous administration of ascorbate can safely result in plasma levels as high as 20 mM, far beyond what is necessary to kill cancer cells^7^. Over the last decade, multiple trials have demonstrated the safety of ascorbic acid alone or in combination with chemotherapy^8^. Some studies suggest that ascorbate could even alleviate the adverse effects of chemotherapy without sacrificing treatment efficacy and improve quality of life for cancer patients^9,10^. Additionally, pharmacological ascorbate has shown *in vivo* efficacy in a variety of cancers as a monotherapy and in combination with existing cancer drugs, including agents known to cause DNA damage, such as cisplatin^11–13^. While there is still uncertainty regarding precisely how ascorbate exerts anticancer effects, one established mechanism suggests high-dose ascorbate generates reactive oxygen species (ROS)^14–16^. The propensity for ascorbate to generate ROS and subsequent DNA damage makes it an intriguing agent to pair with drugs that inhibit DNA damage repair, such as Poly(ADP-ribose) Polymerase (PARP) inhibitors which are FDA approved for a subtype of PCa patients whose tumors are defective in homologous recombination DNA repair.

PARP refers to a family of proteins that provide diverse enzymatic functions^17,18^. PARP-1 is the most abundant member of the PARP family and is intimately involved in the base-excision repair (BER) process of DNA repair^19^. BER is activated in response to DNA damage and enables repair of single-stranded breaks (SSB). Inhibition of PARP-1 compromises BER and causes accumulation of SSB which subsequently become double-stranded breaks (DSB) during DNA replication. Briefly, PARP inhibitors (PARPi) function as NAD+ mimetics that block the active site of PARP-1 and PARP-2, preventing the signaling cascade needed to repair SSBs. DSB are preferentially repaired by the high-fidelity homologous recombination (HR) pathway which includes two well-known tumor suppressor proteins *BRCA1* and *BRCA2*. Accumulation of DNA damage leads to genomic instability and, ultimately, cell death^20–22^. The potential for PARP inhibitors to selectively kill cells with known deficiencies in DNA damage repair (DDR) pathways has led to FDA approval in certain tumor types harboring defects in DNA damage repair, including PCa.

Two recent Phase III trials demonstrated efficacy in combining abiraterone with PARP inhibitors (MAGNITUDE: NCT0374861, niraparib; PROpel^23^: NCT03732820, Olaparib). MAGNITUDE was preselected for HR defects, whereas PROpel found that there were responses to the combination of PARP inhibition and abiraterone irrespective of HR status^23^. While less than 5% of primary prostate cancers harbor HR defects^24^, these defects are elevated in mCRPC, albeit only to ~23%^25^. While FDA approved for mCRPC and having shown benefit in combination with abiraterone, it is critical to combine PARP inhibition with other agents in order to expand the cohort of patients who might benefit from PARP inhibitor therapy. Ultimately, the combination of ascorbate, which has demonstrated tolerability and efficacy in killing cancer cells by generating DNA-damaging ROS, and PARP inhibitors, which are known to impair a cancer cell’s ability to repair DNA damage and have already demonstrated efficacy in treating advanced PCa as a monotherapy, could provide a new treatment modality for the mCRPC population.

We hypothesized that DNA-damaging ROS generated by ascorbate would pair well with PARP inhibitors and their ability to impede DNA damage repair. Combination studies demonstrated a decrease in CRPC proliferation *in vitro* and *in vivo*. The combination of olaparib and ascorbate was so potent as to demonstrate synergy in two distinct *in vitro* models. Mechanistic studies showed a significant increase in ROS accumulation and DNA damage supporting the initial hypothesis. These results suggest the combination of ascorbate and PARP inhibition could be an effective treatment in mCRPC.

## Results

### Ascorbate alters energy dynamics of CRPC cells and generates ROS resulting in DNA damage

Ascorbate toxicity has been demonstrated in several different types of cancer cells; however, the evidence supporting its use in advanced PCa is minimal^8–13^. It has been well-established that pharmacological ascorbate generates DNA-damaging ROS; however, recent data in pancreatic cancer models also suggest that ascorbate can alter cellular bioenergetics^26^. No studies exploring bioenergetics have been performed in PCa. The anti-tumor effects of ascorbate were impacted by both iron and pyruvate concentration in the culture media (**Supplementary Figure 1A**), but manipulating these culture media parameters did not impact tumor cell growth in the absence of ascorbate (**Supplementary Figure 1B and Supplementary Figure 1A**). This study will be the first to identify the novel role of ascorbate enhancing PCa therapeutics.

To assess the potential anti-tumor effects of ascorbate, C4-2 cells conditioned as described in Materials & Methods as well as 22Rv1 cells were cultured in DMEM and treated with 1 mM ascorbate for up to 6 hours. The extracellular concentration of pyruvate was assayed at several time points during treatment. Within thirty minutes of initiating treatment with ascorbate, the concentration of pyruvate decreased by approximately 50% for C4-2 cells compared to untreated control (p=0.020; **Figure 1A, top**). Similarly, within four hours of treatment with ascorbate, 22Rv1 cells saw a greater than 20% reduction in pyruvate concentration (p=0.048, **Figure 1A, bottom**) compared to untreated control. Additionally, within two hours of ascorbate administration, the amount of intracellular ATP decreased by over 50% (p=0.049; Figure 1A, top) or 80% in C4-2 and 22Rv1 cells (p=7.83e-4; Figure 1A, bottom), respectively. These findings were associated with a 55% decrease in the percentage of surviving C4-2 cells within four hours compared to untreated control (p=0.010; **Figure 1A, top**) and by nearly 50% decrease in surviving 22Rv1 cells within 6 hours (p=0.011, **Figure 1A, bottom**). These findings indicate the anti-tumor effects of ascorbate are associated with altered cellular metabolism that precludes reduction in cancer cell viability.

**Figure 1.**
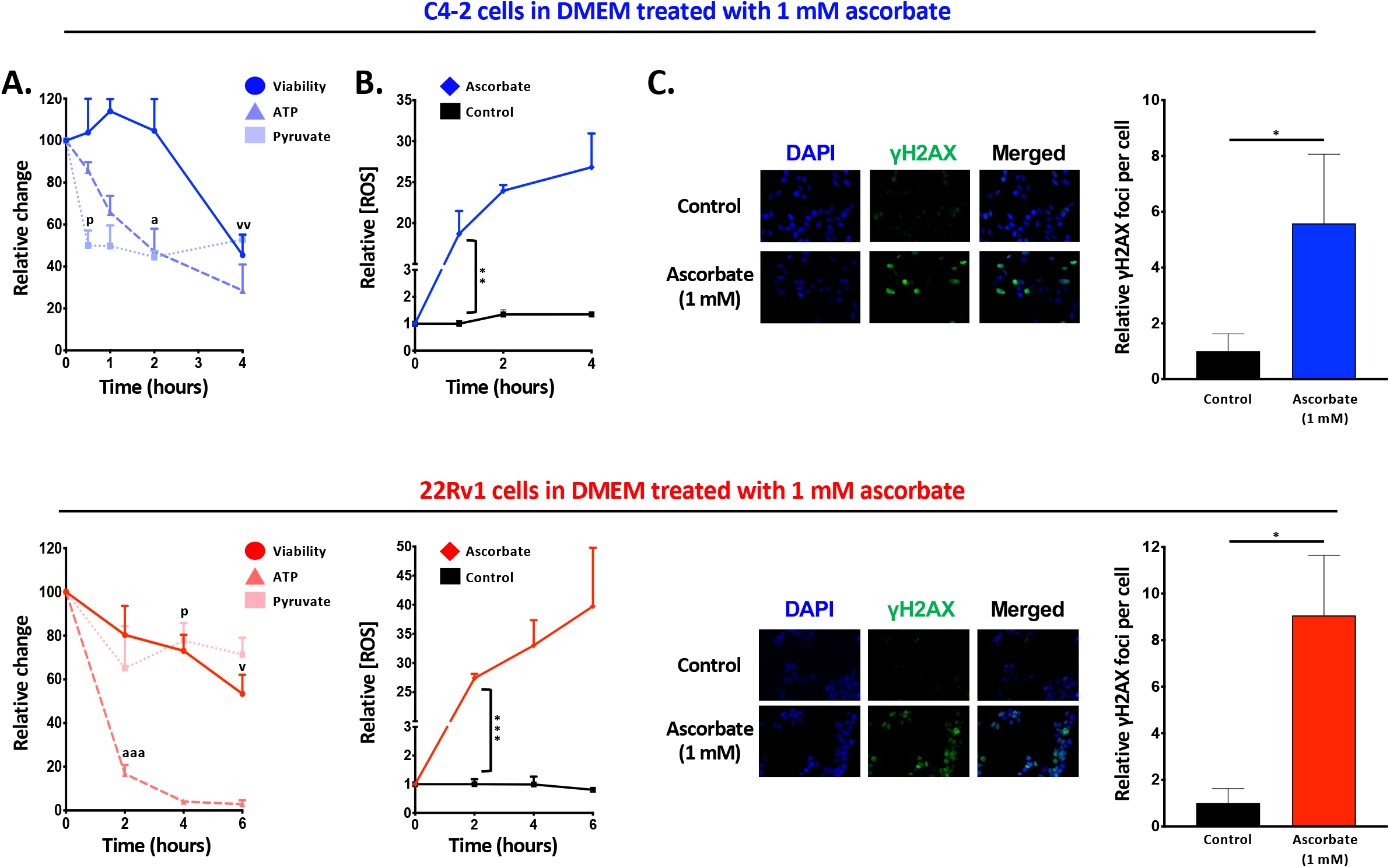
Ascorbate alters energy mechanics of CRPC cells and generates ROS resulting in DNA damage. A) C4-2 cells (top) were conditioned in DMEM, seeded onto 96-well plates and allowed to settle overnight. Wells were treated with 1 mM ascorbate, after which levels of extracellular pyruvate (square), intracellular ATP (triangle) and cell viability (circle) were assessed at the indicated time points. 22Rv1 cells (bottom) were plated in DMEM, treated, and analyzed as described for C4-2 cells. B) C4-2 cells (top) and 22Rv1 (bottom) were plated as described in (A), and levels of ROS were assessed at the indicated time points. C) C4-2 cells (top) and 22Rv1 cells (bottom) were plated as described in (A). C4-2 cells (2 hours) and 22Rv1 cells (4 hours) were treated with 1 mM ascorbate and assessed for formation of γH2AX foci as a surrogate marker for DNA damage. Data are depicted as mean relative to vehicle control ± SEM of at least three independent biological replicates. Statistical significance was determined by Student’s t-test. * denotes P<0.05 compared to untreated control, ** denotes P<0.01 compared to untreated control, *** denotes P<.001 compared to untreated control, *a* denotes P<0.01 compared to ATP levels at 0 hr treatment, *aaa* denotes p<.001 compared to ATP levels at 0 hr treatment, *p* denotes P<0.05 compared to pyruvate levels at 0 hr treatment, *v* denotes P<0.05 compared to viability at 0 hr treatment, *vv* denotes P<0.01 compared to viability at 0 hr treatment.

The combined import of these data is to establish that treatment with high-dose ascorbate affects CRPC cell energy mechanics as evidenced by a decrease in extracellular pyruvate, intracellular ATP and ultimately cell viability. These observations are explicable by considering ascorbate-induced ROS and the effects of ROS on cancer cell metabolism. Cancer cells have constitutively higher levels of pro-tumorigenic ROS^27^ than non-transformed cells; however, excess ROS can damage mitochondrial DNA as well as key proteins in the electron transport chain necessary for generating ATP by oxidative phosphorylation^28^. Additionally, an excess of ROS forces cancer cells to shift substrates from the ATP-generating Krebs cycle into the Pentose Phosphate pathway which generates NADPH, a key antioxidant^29^. Thus there is a dual mechanism involving the impairment of oxidative phosphorylation and glycolysis by which ascorbate-mediated ROS could rapidly deplete cellular ATP. To determine whether elevated ROS was associated with the observed altered cancer cell metabolism and subsequent reduction in cancer cell viability, it was necessary to establish the timing and degree to which ascorbate generates ROS in CRPC models.

A time-dependent accumulation of ROS was observed after 60 minutes of ascorbate treatment with a greater than 18-fold increase in measured ROS for C4-2 cells treated with ascorbate compared to untreated control (p=0.008, **Figure 1B, top**) and a 27-fold increase observed in 22Rv1 cells (p=2.64×10^-4^, **Figure 1B, bottom**) within 2 hours of ascorbate treatment. Immunofluorescent staining for yH2AX showed a 5.5-fold increase in yH2AX foci in C4-2 cells treated with ascorbate for two hours compared to untreated control (p=0.031, **Figure 1C, top**) and a 9-fold increase in 22Rv1 cells treated with ascorbate for four hours (p=0.027, **Figure 1C, bottom**).

These data suggest that treatment with high-dose ascorbate generates cytotoxic levels of ROS that can impair cell metabolism and generate DNA damage. Whether these mechanisms are independent of each other or related is unclear; however, these data indicate that ascorbate can kill CRPC cells making it an attractive therapeutic option and a candidate for combination with other treatments with varying mechanisms of action. Given the observed increase in DNA damage associated with ascorbate monotherapy, it was hypothesized that therapeutic benefit could be achieved by adding an agent which prevents the repair of DNA damage, such as a PARP inhibitor.

### Treatment with ascorbate and Olaparib synergistically inhibits CRPC cell proliferation *in vitro* by generating DNA damage

Three different PARP inhibitors, Olaparib (**Figure 2A, left**), Niraparib (**Figure 2A, middle**) and Talazoparib (**Figure 2A, right**) were assessed for efficacy in treating C4-2 cells (**Figure 2A, top**) and 22Rv1 cells (**Figure 2A, bottom**). All three PARP inhibitors demonstrated a dosedependent ability to inhibit CRPC cell proliferation as monotherapy which was not impacted by pyruvate concentration (**Supplementary Figure 2B**). Intriguingly, the addition of a non-lethal dose of ascorbate significantly augmented the toxicity of all three PARP inhibitors in both CRPC models. Given these promising results, whether the interaction between ascorbate and PARP inhibitors could be considered synergistic was determined. Olaparib, being the most clinically advanced of the three PARP inhibitors tested, was used to generate additional growth curves with ascorbate. Data were analyzed using the program CompuSyn to develop a combination index for the combination of Olaparib and ascorbate in both C4-2 cells (**Figure 2B, top**) and 22Rv1 cells (**Figure 2B, bottom**). For both CRPC models, the majority of data points generated fall into the region where the combination index is less than one, suggesting a strongly synergistic relationship between Olaparib and ascorbate *in vitro*.

**Figure 2.**
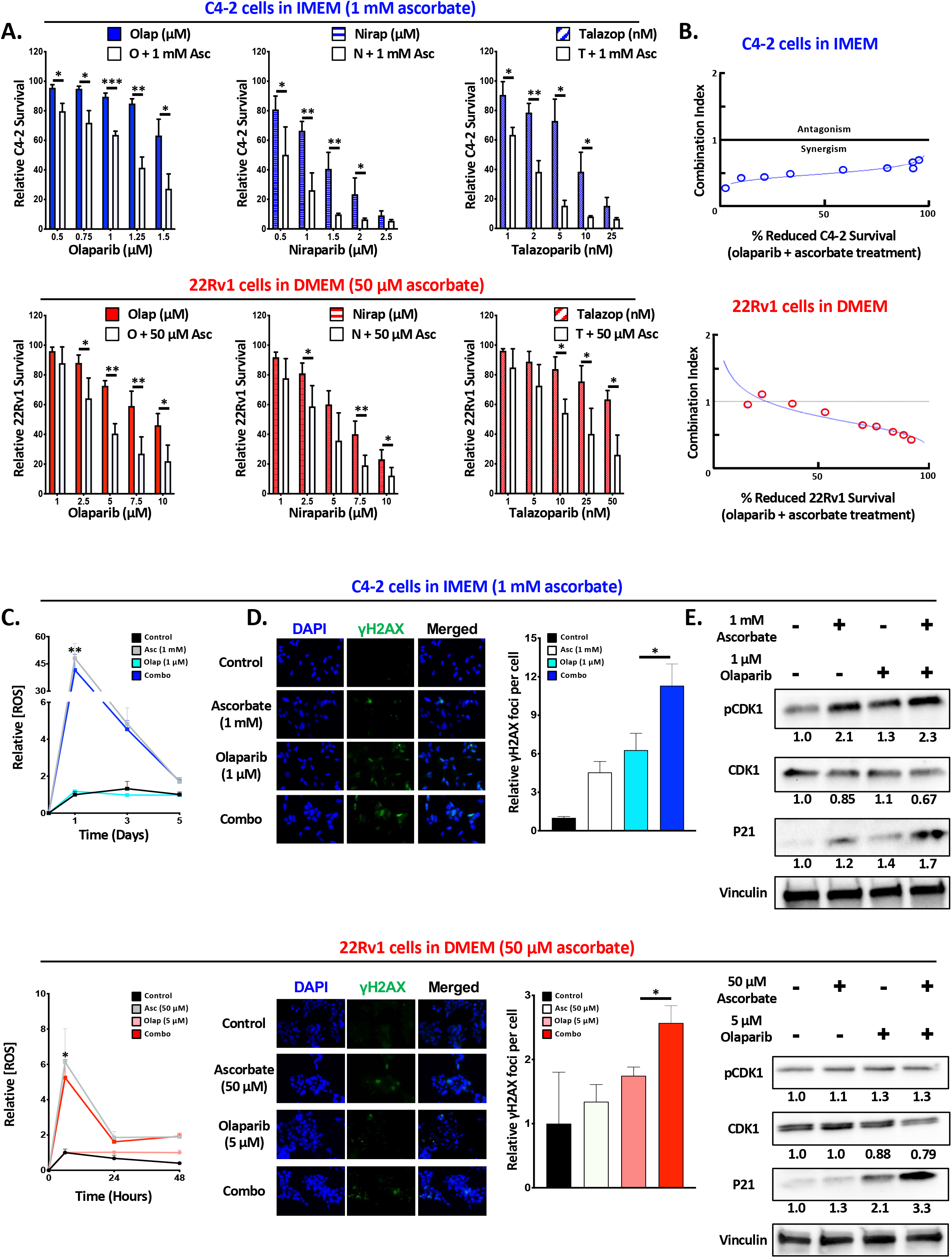
The combination of ascorbate and olaparib synergistically inhibits CRPC cell proliferation *in vitro* by generating DNA damage. A) Equivalent densities of C4-2 cells (top) and 22Rv1 cells (bottom) were seeded on 96-well plates in IMEM and DMEM, respectively. Cells were treated with a combination of ascorbate and either olaparib (left), niraparib (center) or talazoparib (right) at the indicated concentration for 5 days. Cell survival was assessed using the Picogreen assay. B) CompuSyn was used to generate a combination index (CI) for the combination of ascorbate and olaparib in C4-2 cells (top) and 22Rv1 cells (bottom). CI > 1 denotes antagonism, CI =1 denotes an additive effect, CI < 1 denotes synergism. C) Equivalent densities of C4-2 cells (top) or 22Rv1 cells (bottom) were seeded on a 96-well plate and treated with the indicated amount of ascorbate and olaparib alone and in combination. The levels of ROS were assayed at the indicated time points. D) Equivalent densities of C4-2 cells (top) or 22Rv1 cells (bottom) were seeded on 6-well plates and treated with the indicated amount of ascorbate and olaparib alone and in combination for 3 days. Cells were assessed for formation of γH2AX foci as a surrogate marker for DNA damage. E) C4-2 cells (top) or 22Rv1 cells (bottom) were seeded on 10 cm^2^ plates and treated with the indicated amount of ascorbate and olaparib alone and in combination for 3 days. Culture media and attached cells were collected for western blot analysis of proteins associated with G2/M checkpoint. Data are depicted as mean relative cell survival (compared to vehicle control) ± SEM of at least three independent biological replicates. Statistical significance was determined by Student’s t-test. * denotes P<0.5, ** denotes P<0.1, *** denotes p<.001.

Given the capacity of ascorbate to generate ROS as a single agent, the impact of PARPi on ascorbate-driven ROS generation was determined. Even the non-lethal dose of ascorbate was associated with significant ROS accumulation in C4-2 cells, 48-fold increase compared to untreated control after 24 hours (p=0.009, **Figure 2C, top**), and 22Rv1 cells, 6-fold increase compared to untreated control after 6 hours (p=0.041, **Figure 2C, bottom**). Of note, the ROS level appeared to peak within 24 hours and rapidly decline, mirroring the rapid metabolism of ascorbate seen in human subjects. Furthermore, the addition of Olaparib did not impact the extent or duration of ROS accumulation.

Utilizing yH2AX as a marker for DNA double strand breaks, C4-2 cells treated with the combination of ascorbate and olaparib generated 11-fold more yH2AX foci relative to untreated control compared to a 6-fold increase by Olaparib alone (p=0.017, **Figure 2D top**). A similar pattern was observed in 22Rv1 cells where a 2.5-fold increase was generated by the combination treatment relative to untreated control compared to 1.7-fold increase by Olaparib alone (p=0.019, **Figure 2D bottom**). Interpreting the ROS and yH2AX data together suggests that combination treatment does not generate additional ROS compared to monotherapy; however, increased accumulation of DNA damage is observed. This is consistent with the hypothesis that PARP inhibition, while having no effect on the presence of ROS, would cause a delay in the repair of DNA damage generated by the presence of ROS. Ultimately, ascorbate-induced ROS generation leads to an elevation in DNA double-strand breaks, and the addition of PARPi to ascorbate results in a further accrual of DNA damage.

To examine the impact of the tested therapies on DNA repair factors, protein expression was assessed with combination therapy in CRPC models. Western blot analysis for C4-2 cells treated with ascorbate and Olaparib alone or in combination showed a significant increase in p21 protein concentration and activity, as evidenced by an increase in phosphorylated CDK1, compared to either monotherapy or untreated control (**Figure 2E, top**). A similar pattern was seen in 22Rv1 cells (**Figure 2E, bottom**). p21 activity is increased in response to DNA damage and one of its many functions is to inhibit CDK1 by phosphorylation to halt the cell cycle at the G2/M checkpoint and allow for repair of DNA damage^30^. Taken together, these data suggest that the combination of ascorbate and Olaparib inhibit CRPC cell proliferation synergistically *in vitro* by generating DNA damage.

### The combination of Olaparib and ascorbate slows tumor growth *in vivo*

Given the promising *in vitro* results generated by combining of ascorbate and Olaparib, the impact of this treatment *in vivo* was next investigated. Xenografts were generated by injecting C4-2 cells into both flanks of two cohorts of immunocompromised mice, castrated and noncastrated. The mice were then randomly assigned to receive daily IP injections of saline, ascorbate alone, Olaparib alone or the combination of Olaparib and ascorbate. Tumor growth was measured three times weekly. Mice treated with the combination of ascorbate and Olaparib showed significantly increased tumor doubling time compared to mice treated with Olaparib alone (p=5.00×10^-4^) or ascorbate alone (p=0.015) in both non-castrated (**Figure 3A, left**) and castrated conditions p=0.007 and p=0.022, respectively, **Figure 3A, right**). Upon reaching the endpoint, tumors were harvested, and tissue underwent IHC staining for p21. Tumors from mice treated with the combination of ascorbate and Olaparib showed significantly increased percent positive cells compared to mice treated with Olaparib alone (p=0.050) or ascorbate alone (p=0.008) in both non-castrated (**Figure 3B, left**) and castrated conditions (p=0.050 and p=0.016, respectively, **Figure 3B, right**) with minimal impact on animal weight (**Supplementary Figure 3**). These data suggest the combination of ascorbate and olaparib slows tumor growth *in vivo*, highlighting a potential novel combinatorial therapeutic regimen to improve patient outcome

**Figure 3.**
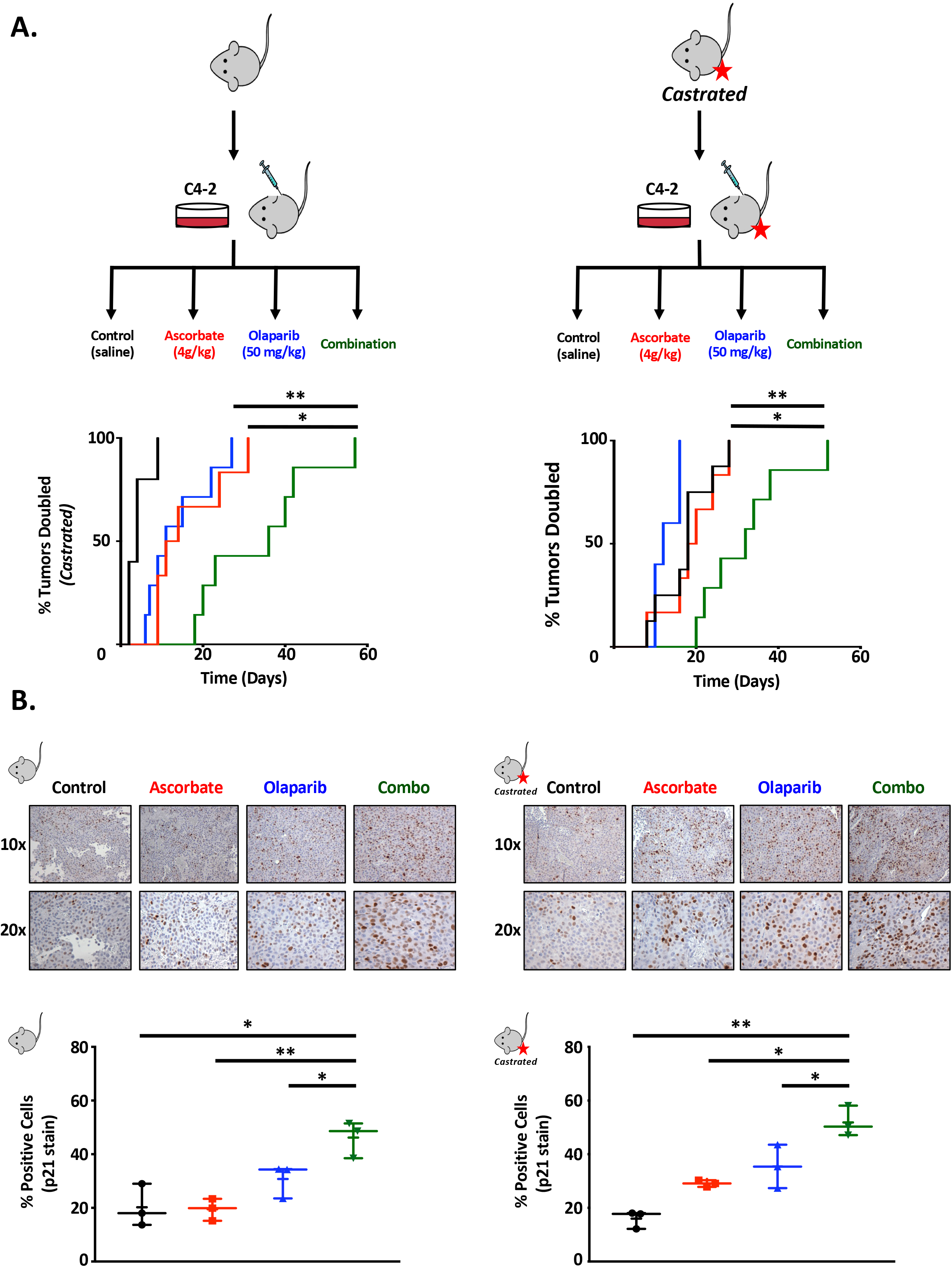
The combination of olaparib and ascorbate slows tumor growth *in vivo*. A) C4-2 xenografts were generated in either non-castrated (left) or castrated (right) NOD/SCID mice. Mice that developed tumors were randomly assigned into cohorts receiving daily IP injections of normal saline, 50 mg/kg olaparib, 4 g/kg ascorbate or a combination of 50 mg/kg olaparib and 4 g/kg ascorbate. Tumor volume was measured by calipers and calculated by V_tumor_=(short distance)^2^ x long distance x 0.5236. Statistical significance was determined by Log-rank (Mantel-Cox) test. B) Harvested tumors underwent IHC analysis for P21. Representative images are shown at 10x and 20x and quantitative analysis is shown below. Three slides, each from a different tumor, were chosen from each treatment group, and five randomly chosen fields from each slide were photographed and scored. Data depicted represent mean % positive cells. Statistical significance was determined by Student’s t-test. * denotes P<0.05, ** denotes P<0.01.

## Discussion

As discussed in the introduction, there is a high prevalence of DDR alterations in PCa, and these are associated with high-grade histology and metastatic disease^31^. Multiple clinical trials have shown that PARP inhibition in DDR defective mCRPC is associated with improved progression free survival (PFS)^32^. Two PARP inhibitors have received FDA approval for use in mCRPC, Olaparib based on an increase in PFS observed in the PROfound trial and Rucaparib based on improved radiographic and prostate specific antigen response in the TRITON-2 trial; similar studies are underway for Talazoparib and Niraparib^33^. A key limiting factor in the use of PARP inhibitors in mCRPC is that some trials indicate necessity of DDR alterations in order to see efficacy, while other trials demonstrate some benefit in non-mutant tumors. While mCRPC is associated with a higher rate of DDR alterations relative to other cancers, such mutations are still only present in a minority of mCRPC cases. For example, in the PROfound trial, only 28% of the screened population had a qualifying mutation^34^. This suggests that broader application of PARP inhibitors will likely require combination with other cancer therapies. Currently, there are multiple clinical trials studying the potential benefits of treating PCa by combining PARP inhibition with various therapies including ADT, immune checkpoint inhibitors, kinase inhibitors, and radiation, with the latter combination attempting to capitalize on the ability of radiation therapy to induce DNA damage^35^.

The therapeutic implication of combining PARP inhibitors with an agent that generates DNA damage was the impetus for choosing ascorbate in the combination studies presented here. There are many DNA damaging agents from which to choose, and ascorbate was selected for several reasons including relative affordability compared to other chemotherapy agents, a favorable tolerability profile, and a growing body of evidence showing efficacy in treating a variety of cancers as monotherapy and in combination with standard of care chemotherapy and radiation treatments^8–13^. The proposed mechanism by which ascorbate and PARP inhibition might prove effective in CRPC was based upon the tendency for ascorbate to generate ROS when dosed at high levels; ROS accumulation generates DNA damage which PARP inhibition would augment and perpetuate leading to CRPC cell death. The data presented here support this mechanism as ascorbate was shown to generate ROS *in vitro* and the combination of PARP inhibition and ascorbate produced a statistically significant increase in DNA damage as measured by γH2AX foci formation *in vitro*. It seems that ascorbate, as a DNA damaging agent, pairs well with PARP inhibitors.

Additional *in vitro* assays using ascorbate as monotherapy demonstrated a time-dependent decrease in pyruvate and ATP levels subsequently followed by a significant decrease in CRPC cell proliferation. These results correlate with *in vitro* data derived from experiments in breast cancer that describe an ascorbate-induced “energy catastrophe” evidenced by rapid depletion of ATP and cell death^36^. While these results were generated in the absence of any PARP inhibition, other studies have produced evidence that, in ATP-deficient situations, PARP enzymes generate ATP needed to properly execute BER^37^. If true, then PARP inhibition could augment the depletion of ATP that results from ascorbate treatment. This suggests the mechanism driving the synergy between these two agents may be more intricate than simply ascorbate inducing DNA damage and PARP inhibition impeding the repair process.

Further literature review sheds light on yet another facet that may influence the interaction between ascorbate and PARP inhibitors. It is well established that ascorbate is an essential cofactor for a variety of enzymes known as dioxygenases. One such family of enzymes, the ten-eleven translocation (TET) methylcytosine dioxygenases, catalyze the hydroxylation of methylated cytosine bases in DNA, one of the initial steps in DNA demethylation. DNA methylation is an important process by which cells can regulate expression of specific genes and much has been written about the significance of such epigenetic regulation in the development, progression, and treatment of cancer in general. Ascorbate’s association with the TET family suggests a role in epigenetic control of the genome and could have significant implications for cancer treatment. A plethora of *in vitro* and *in vivo* studies performed in a wide range of solid and hematological malignancies have consistently demonstrated that treatment with ascorbate, either as monotherapy or in combination with established epigenetic modifiers, significantly increases TET catalytic activity^38^. The tendency for PARP inhibitors to form complexes by directly binding PARP enzymes at sites of DNA damage makes them an intriguing target for combination with epigenetic modifiers^39^. At least one set of *in vitro* and *in vivo* experiments in ovarian cancer demonstrated that the combination of Olaparib and one such epigenetic modifier, 5-azacitidine, showed significant anticancer effect^40^. This suggests that ascorbate’s role in potentially augmenting demethylation of DNA could underlie yet another mechanism driving the tantalizing anticancer effects seen in combination with PARP inhibition.

The data presented here contribute to a rapidly growing body of evidence that ascorbate could be an effective weapon in the war against cancer either alone or in combination with other agents; however, there are hurdles yet to clear, namely the lack of high-quality data from randomized controlled trials. According to clinicaltrials.gov, there are only five actively recruiting trials examining pharmacological ascorbate in cancer, all of which are categorized as Phase 2. The limited data that exist in human subjects are promising, and there is even one study that has looked at PARP inhibition and ascorbate in humans: eight patients with a variety of stage IV malignancies were treated with either Olaparib, Niraparib, or Talazoparib and pharmacological ascorbate; five patients showed partial response and three showed complete response with no grade three toxicity reported^41^. Results presented herein were used as preclinical data to form the basis for a clinical trial (NCT05501548). The trial schema is presented in **Figure 4**.

**Figure 4.**
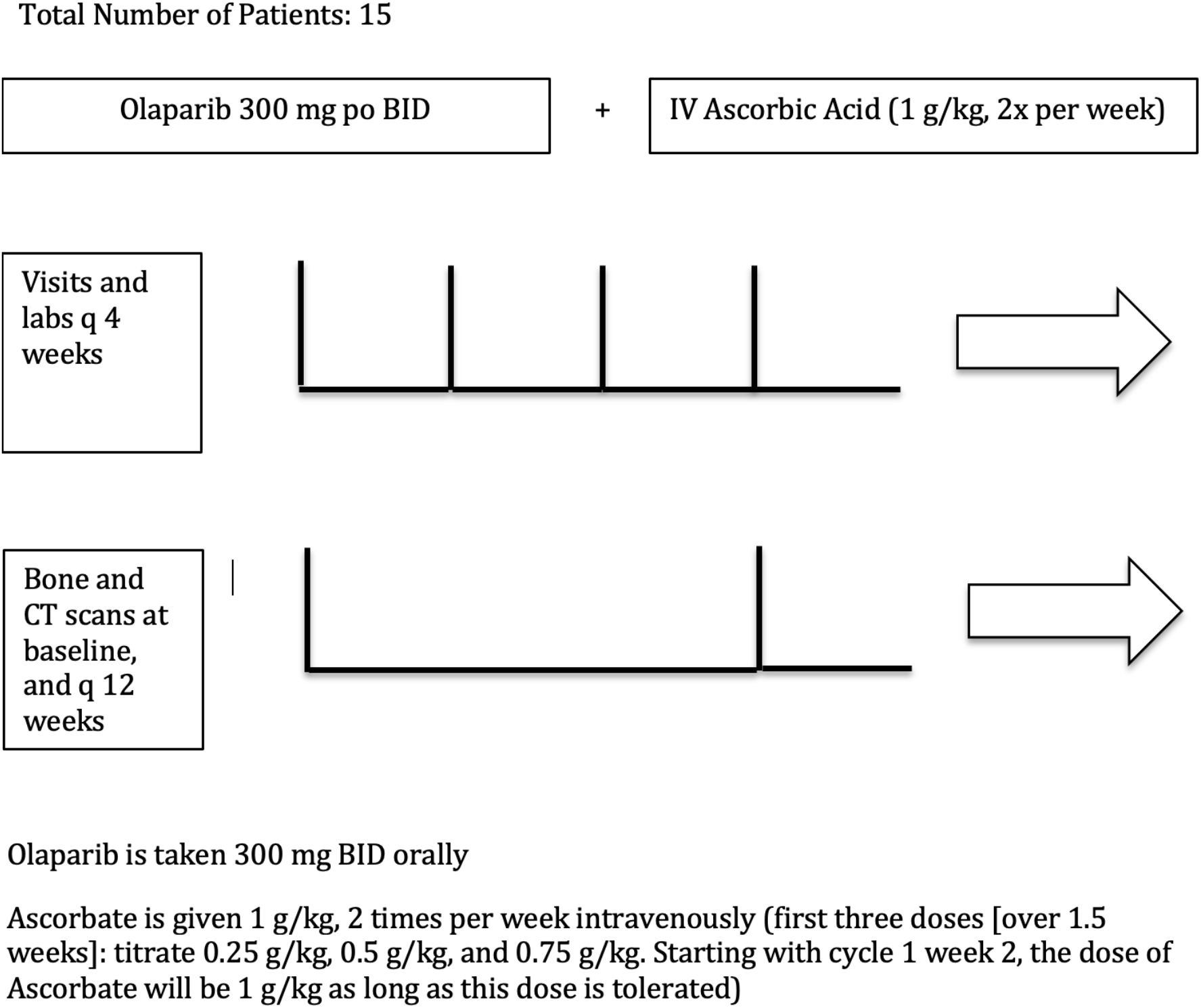
Trial schema for NCT05501548: Phase II Study of PARP Inhibitor Olaparib and IV Ascorbate in Castration Resistant Prostate Cancer.

In summary, this investigation produced molecular and translational evidence that the combination of ascorbate and Olaparib could be an effective treatment in mCRPC. Currently, the most evidence-based mechanism is based on ascorbate generating DNA-damaging ROS and PARP inhibitors preventing the repair of that damage. However, additional evidence suggests a potentially multifaceted mechanism driving the synergy observed between ascorbate and PARP inhibition based on ascorbate altering cellular energy mechanics and playing a role in epigenetic regulation of the genome. The depth and breadth of these proposed interactions suggest broad applicability and less potential for resistance to occur in response to treatment with ascorbate and PARP inhibition. Currently, there is a dearth of high-quality data in human subjects, and an emphasis must be placed on performing large-scale randomized controlled trials as soon as possible. This investigation has produced compelling evidence for the use of ascorbate and PARP inhibition in CRPC, while also contributing to the overall assertion that ascorbate should be considered as an anticancer agent in general.

## Materials and Methods

### Cell Lines, cell culture and treatment

C4-2 and 22Rv1 cells were purchased from ATCC, authenticated by ATCC, and assayed for mycoplasma upon thawing. C4-2 cells are categorized as p53-functional, express a mutated version of the androgen receptor (AR), and are therefore capable of proliferating in androgen deprived conditions. 22Rv1 cells are heterozygous for p53 mutation, express a splice variant of AR known as AR-V7, and are also capable of proliferating in androgen-deprived conditions. C4-2 cells were cultured and maintained in Improved Minimum Essential Medium (IMEM) (Thermo Fisher Scientific, 10024CV) supplemented with 5% FBS (fetal bovine serum, heat inactivated), 1% L-glutamine (2 mmol/l), and 1% penicillin-streptomycin (100 units/ml). 22Rv1s cell were cultured and maintained in Dulbecco’s Modified Eagle Medium (DMEM) (Thermo Fisher Scientific, 10017CV) supplemented with 10% FBS, 1% L-glutamine (2 mmol/l), and 1% penicillinstreptomycin (100 units/ml). All cells were cultured at 37°C with 5% CO2. For indicated experiments, the cell culture media was supplemented with sodium pyruvate (Sigma-Aldrich P2256-5G made at 100 mM stock stored at 4^°^C), iron nitrate (Sigma-Aldrich 216828-100G made at 50 mM stock stored at 4^°^C), or iron chloride (Sigma-Aldrich 236489-5G made at 50 mM stock stored at 4^°^C). Treatment then proceeded with Sodium L-ascorbate (Sigma Aldrich A4034-100G made at 100 mM stock and stored at 4^°^C), olaparib (Selleck S1060 made at 100 mM stock and stored at −20^°^C), niraparib (AdooQ BioScience A11026 made at 100 mM stock and stored at −20^°^C), and/or talazoparib (Selleck S7048 made at 10 mM stock and stored at −80^°^C) depending on the experiment. For experiments in which cells were plated and treated in the alternate cell culture medium (C4-2 in DMEM and 22Rv1 in IMEM), cells were initially thawed into their original medium and passaged 3 times in the alternate medium prior to “condition” them prior to use for experiments.

### Proliferation Assays

Equivalent densities of C4-2 or 22Rv1 cells were seeded in their respective media on clear, Poly-L-lysine coated 96-well plates and allowed to adhere overnight. Cells were then pre-treated and or treated with the reagents previously described. After treatment, the media was discarded, cells were gently washed in PBS several times and cells were lysed in 100 uL of dH2O for 1 hour at 37^°^C. Cells were then incubated with the Quanti-IT Pico Green dsDNA reagent (Invitrogen P7581) per manufacturer’s instructions. Data were collected using a BioTek Synergy HT plate reader. Cells were compared to Day 0 for normalization.

### Metabolic Assays

#### Pyruvate Assay

Equivalent densities of C4-2 or 22Rv1 cells were seeded on Poly-L-lysine coated 96-well plates and allowed to adhere overnight. At indicated time points during treatment, cell culture media was collected and pyruvate concentration was assessed using the Pyruvate Colorimetric Assay Kit (BioVision K609-100) per manufacturer’s instructions. Absorbance was detected using a BioTek Synergy HT plate reader.

#### ATP Assay

Equivalent densities of C4-2 or 22Rv1 cells were seeded on Poly-L-lysine coated, white-walled 96-well plates (Sigma-Aldrich CLS3610-48EA) and allowed to adhere overnight. At indicated time points during treatment, intracellular ATP levels were assessed using the CellTiter-Glo 2.0 Cell Viability Assay (Promega G9242) per manufacturer’s recommendations. Luminescence was detected using a BioTek Synergy HT plate reader.

#### ROS Assay

Equivalent densities of C4-2 or 22Rv1 cells were seeded in their respective media on Poly-L-lysine coated, white-walled 96-well plates (Sigma-Aldrich CLS3610-48EA) and allowed to adhere overnight. Cells were then pre-treated and or treated with the reagents previously described. After treatment, ROS levels were assessed using the ROS-Glo H2O2 Assay (Promega G8821) per manufacturer’s instructions. Briefly, cells were pre-incubated with the ROS substrate for 2 hours at 37^°^C prior to analysis at which point the detection reagent was added for 20 minutes at room temperature. ROS levels were detected by quantifying luminescence using a BioTek Synergy HT plate reader.

### Immunofluorescence

Immunofluorescence (IF) experiments were performed as detailed previously^42^. Briefly, equivalent densities of C4-2 or 22Rv1 cells were seeded on poly-lysine coated coverslips in 6-well plates and allowed to adhere overnight. After treatment, coverslips were washed gently in PBS and fixed for 20 minutes with 3.7% Formaldehyde at room temperature. Cells were stained using γH2AX phospho-S139 (EMD Millipore 16-202A) at 1:500 dilution. Foci were imaged utilizing Dr. Elda Grabocka’s Zeiss Cell Discoverer Confocal Microscope at 40X magnification with at least 5 fields for each replicate. Fiji image software was utilized to quantify foci formation per cell and compared to control samples.

### Immunoblotting

C4-2 and 22Rv1 cells were plated at equal densities in their respective media on Poly-L-lysine coated 10 cm^2^ plates. Generation of cell lysates was described previously^43^. Briefly, 40-50 μg of lysate was resolved by SDS-PAGE, transferred to nitrile membrane and analyzed using the following antibodies: P21 (1:1000, Abcam ab109520), phospho-CDC2 (Tyr15) (1:1500, Cell Signaling Technology 9111S), CDC2 p34 (1:200, Santa Cruz Biotechnology Sc-54) and Vinculin (1:1000, Sigma-Aldrich V9264-200uL)

### Generation of Xenografts

C4-2 cells were cultured and lifted from plates by trypsinization then re-suspended in 100 mL of 50% Matrigel (BD Biosciences) and saline mixture followed by subcutaneous injection in two separate groups of castrated and non-castrated athymic nude mice (age at least 6 weeks old). Once tumors reached 100 mm^3^ in size, mice were triaged into four separate treatment groups and given daily IP injections of either vehicle control (0.09% saline), ascorbate (4g/kg), olaparib (50mg/kg), or a combination of both. Tumor size was measured using calipers every other day. When tumor sizes reached 1000 mm^3^ in size mice were euthanized and their tumors harvested and fixed in 4% formalin in preparation for sectioning. All animal work was done in accordance with IACUC at Jefferson/SKCC.

### IHC

For histological analysis from xenograft tissue, FFPE sections were stained with p21 (Cell signaling S947S) using standard techniques previously described^43^.

### Statistical analysis

All experiments were performed in technical triplicate with at least 3 biological replicates per condition. Data are displayed as mean +/− standard error of the mean (SEM). Statistical significance (p < 0.05) was determined using Student’s t-test.

## Acknowledgements

The authors would like to thank the Schiewer, Shafi, and Knudsen labs for advice and feedback. MJ Schiewer would like to thank the Departments of Urology and Pharmacology, Physiology, & Cancer Biology, as well as the Sidney Kimmel Cancer Center ongoing support. The authors would like to acknowledge the Translational Research/Pathology Core and the Animal Research Facilities at the Sidney Kimmel Cancer Center at Thomas Jefferson University. This research was supported by Pennsylvania Department of Health CURE Grant (to M.J. Schiewer), a Philadelphia Prostate Cancer Biome grant (to M.J. Schiewer), a Prostate Cancer Foundation Young Investigator Award (to M.J. Schiewer), a Prostate Cancer Foundation Challenge Award (to K.E. Knudsen) and the Sidney Kimmel Cancer Center Support Grant NIH P30 CA056036.

---

**Supplementary Figure 1.**
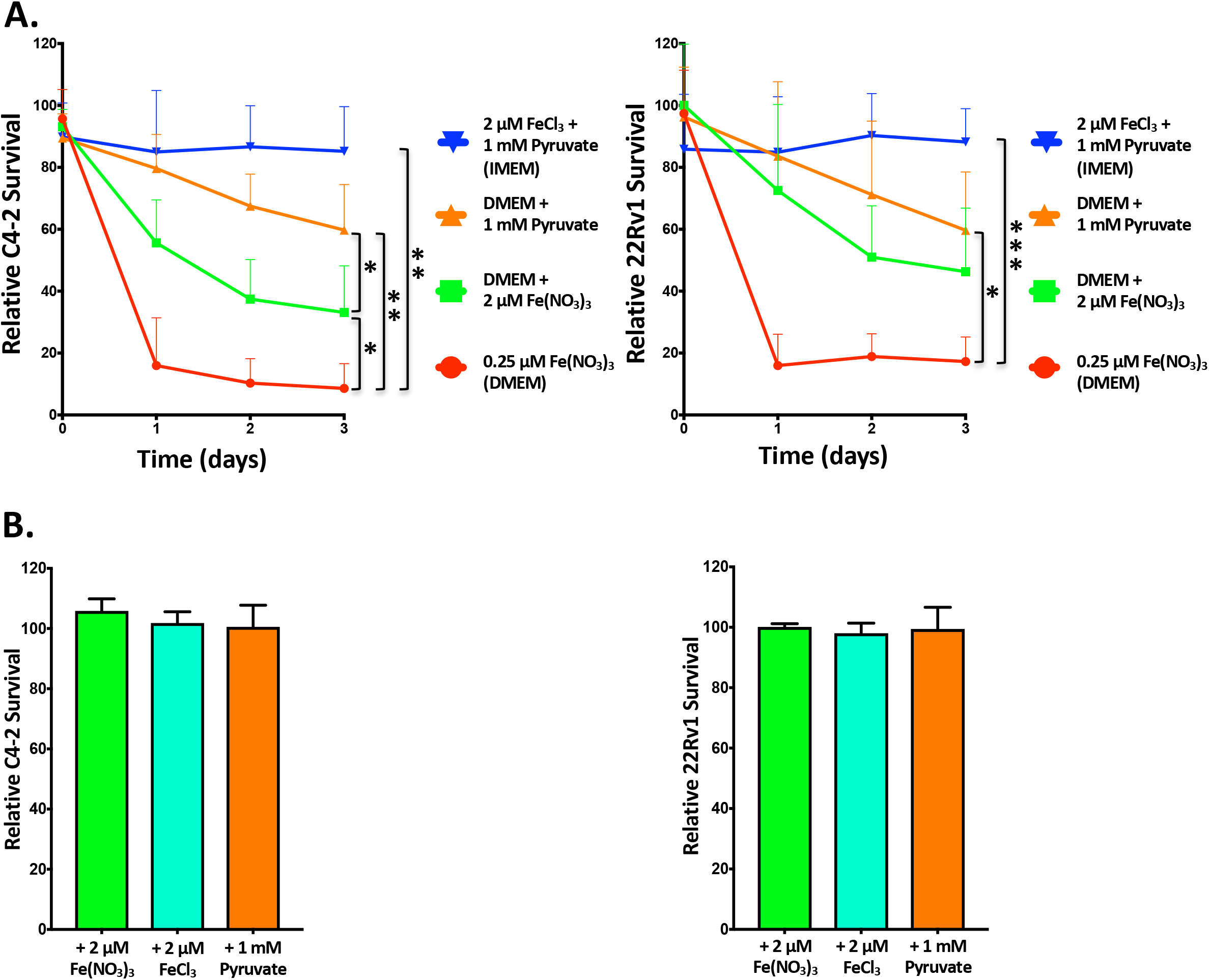
Manipulating [iron] and [pyruvate] has a significant effect on ascorbate toxicity *in vitro*. A) C4-2 (left) or 22Rv1 (right) cells were seeded in DMEM at equal density and allowed to adhere overnight. Sodium pyruvate or iron nitrate was administered, such that the total concentration reached 1 mM for sodium pyruvate or 2 μM for iron nitrate. After 24 hours, cells were treated with 1 mM ascorbate; additional sodium pyruvate or iron nitrate was co-administered in order to maintain the appropriate concentrations. DNA content was quantified using the PicoGreen assay at indicated time points as a means to quantify cell survival. Data are depicted as mean relative cell survival (compared to vehicle control) mean ± SEM of at least three independent biological replicates. B) C4-2 cells (left) or 22Rv1 cells (right) were seeded in DMEM on 96-well plates and allowed to settle overnight. Either sodium pyruvate, iron nitrate or iron chloride was administered to designated wells such that the total concentration reached 1 mM for sodium pyruvate or 2 μM for iron nitrate or iron chloride. After 72 hours, DNA content was quantified using the PicoGreen assay. Statistical significance was determined by Student’s t-test. * denotes P<0.5, ** denotes P<0.1., *** denotes p<.001.

**Supplementary Figure 2.**
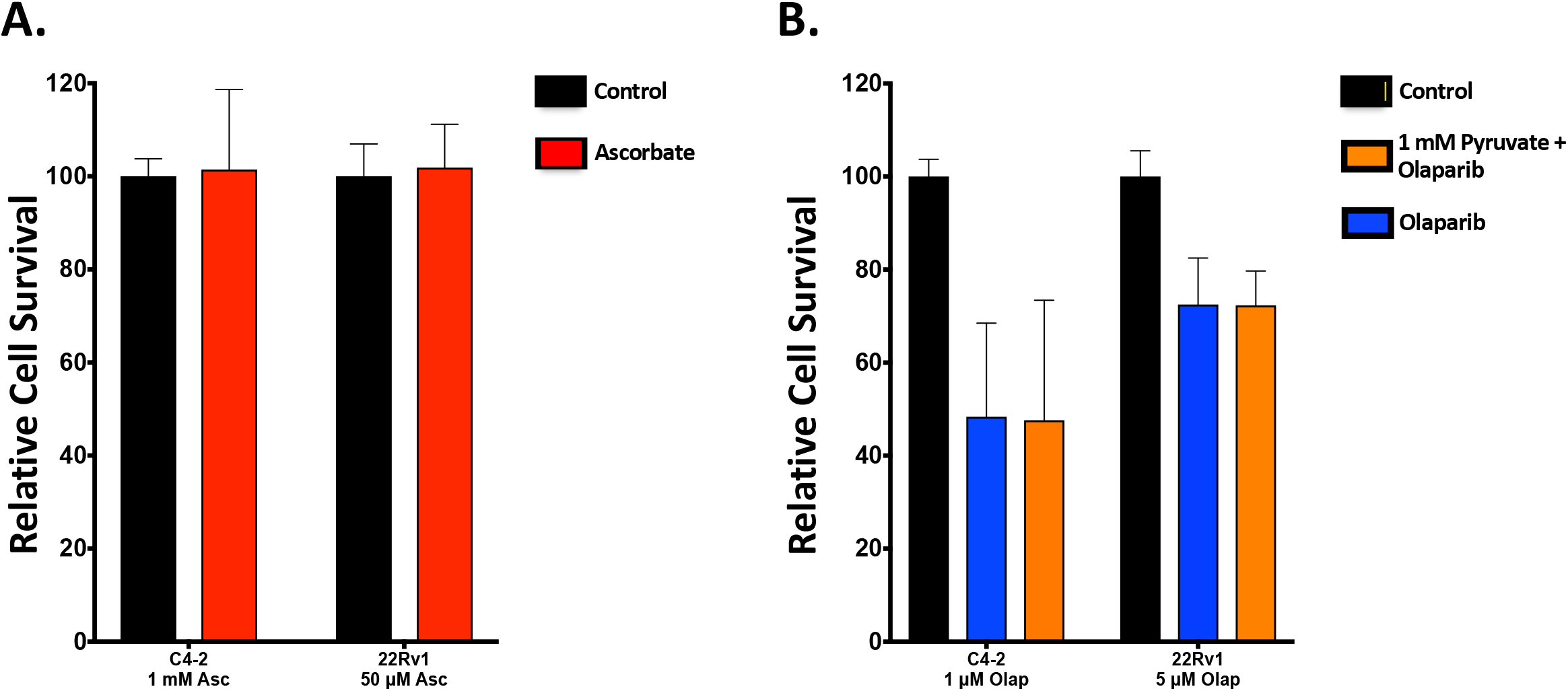
A) C4-2 cells (IMEM) and 22Rv1 cells (DMEM) were seeded on 96-well plates and treated with 1 mM ascorbate or 50 μM ascorbate, respectively for 5 days. Cell survival was assessed using the PicoGreen assay. B) Conditioned C4-2 cells and 22Rv1 cells were seeded in DMEM on 96-well plates and pre-treated with either 1 mM pyruvate or plain DMEM. Cells were treated with 1 μM olaparib (C4-2 cells) or 5 μM olaparib (22Rv1 cells) for 5 days. Cell proliferation was assessed using the PicoGreen assay. Data are depicted as mean relative cell proliferation (compared to vehicle control) ± SEM of at least three independent biological replicates.

**Supplementary Figure 3.**
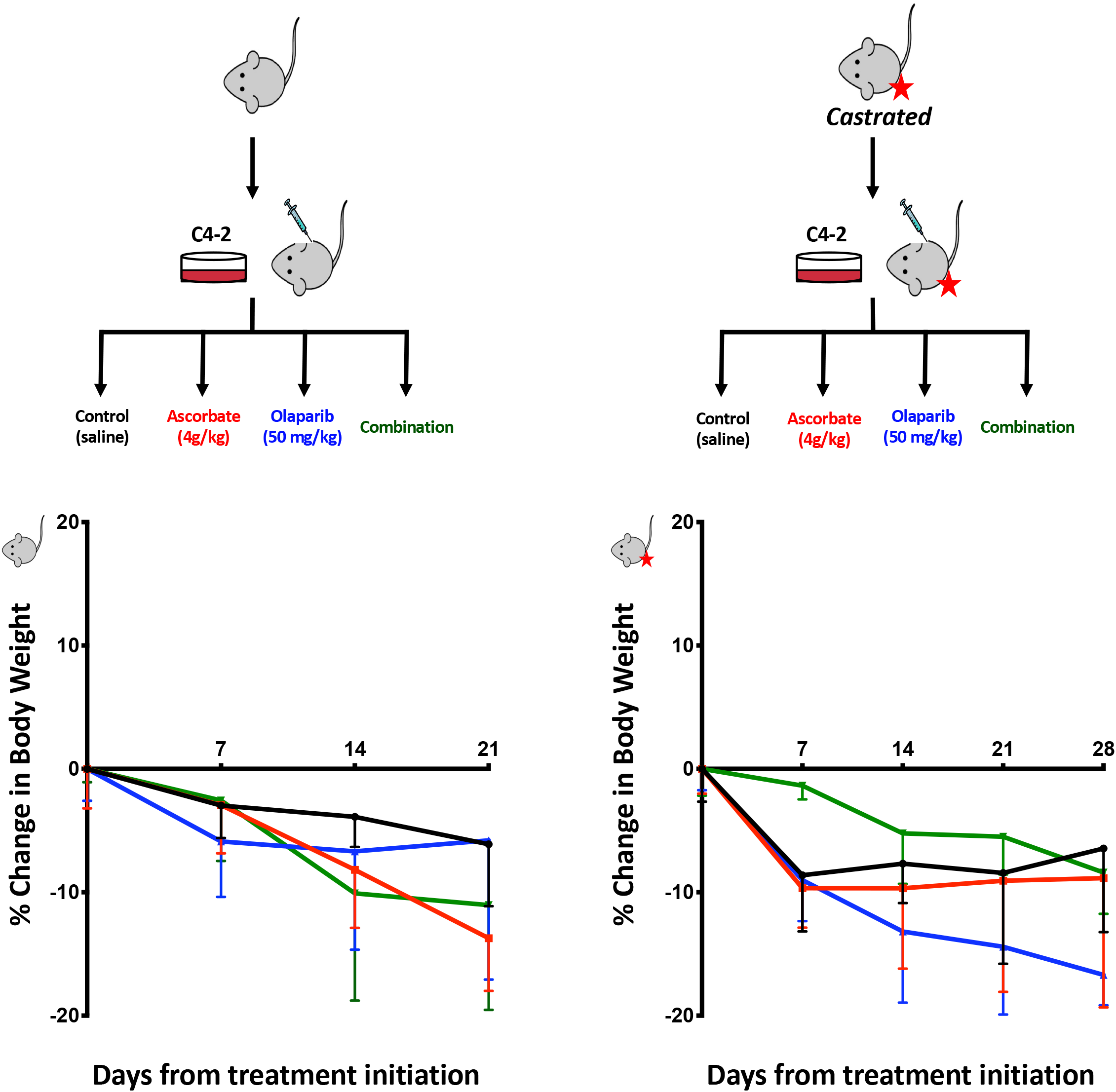
C4-2 xenografts were generated in either non-castrated (left) or castrated (right) NOD/SCID mice. Mice that developed tumors were randomly assigned into cohorts receiving daily IP injections of normal saline, 50 mg/kg olaparib, 4 g/kg ascorbate or a combination of olaparib and ascorbate. Tumor volume was measured by calipers and calculated by V_tumor_=(short distance)^2^ x long distance x 0.5236. Mice were weighed once per week to adjust treatment dosing and monitor for toxicity. Data points represent the average of at least three mice per condition.

